# Quantifying a trait-mediated indirect effect of an environmental stressor on predator-prey dynamics

**DOI:** 10.1101/2025.09.04.674171

**Authors:** Shota Shibasaki, Masato Yamamichi

## Abstract

Trait-mediated indirect interactions have been intensively studied to understand ecological dynamics in biological communities. Yet, little is known about the relative importance of trait-mediated indirect effects of environmental stressors on population dynamics. How does an environmental stressor indirectly affect the population dynamics of a focal species by altering the traits of interacting species and the strength of interspecific interactions? Here, we quantified the direct and trait-mediated indirect effects of an environmental stressor by combining rotifer-alga experiments and Bayesian parameter estimation of a dynamic model. These days, human activities salinize freshwater lakes globally, thereby increasing the death rates and decreasing birth rates of plankton species. Salinity stress is also known to induce cell clumping in certain phytoplankton species. As larger clumps work as a defense trait against gape-limited predation by zooplankton, the salinity stress can affect zooplankton not only directly but also indirectly through phytoplankton trait changes. We first show that a green alga, *Chlamydomonas sphaeroides*, formed larger clumps than the two model species of green algae (*Chlamydomonas reinhardtii* and *Chlorella vulgaris*) under a moderate salinity stress (0.06M NaCl). Then, by tracking the clump size distributions of *Chlamydomonas sphaeroides*, we confirmed that small clumps are more vulnerable to predation by rotifers, *Brachionus calyciflorus*, as previous studies demonstrated. Finally, we co-cultured the green algae and rotifers for a week with and without salinity stress and fitted the Lotka-Volterra predator-prey model. We first estimated how salinity stress increased the rotifer mortality rate using *Chlorella vulgaris*, which seldom showed clump formation. Then, by using *Chlamydomonas sphaeroides*, we estimated how salinity stress increased the mortality rate and decreased the attack rate due to clump formation. We found that salinity stress increased the rotifer mortality rate by more than seventeen-fold, and decreased the attack rate on *Chlamydomonas sphaeroides* to approximately half of that without salinity stress. These results indicate that salinity stress can weaken the predator-prey interaction, and thus salinization can harm freshwater zooplankton species through increasing the mortality rates and decreasing the attack rates. This will be an important step for a quantitative understanding of how environmental stressors can affect community dynamics via trait modifications.

## Introduction

Classic studies in ecology have tended to focus on interspecific interactions mediated by population densities, but recent studies have demonstrated the importance of considering trait dynamics driven by behavioral changes, phenotypic plasticity, or rapid evolution (Bolker et al. 2003, Miner et al. 2005, Bolnick et al. 2011, Hendry 2017). Particularly, trait-mediated indirect interactions (TMII) have been intensively studied to understand community dynamics (Werner and Peacor 2003, Schmitz et al. 2004, Utsumi et al. 2010, Ohgushi et al. 2012, Golubski et al. 2016). For example, trophic cascades were traditionally investigated through the perspective of density-mediated indirect interactions: carnivores reduce the population density of herbivores, which increases plant biomass (Hairston et al. 1960, Estes and Palmisano 1974). Studies on TMII, in contrast, emphasized the importance of behavioral change of intermediary species with the foraging-predation risk trade-off: carnivores may change the behavior of herbivores (e.g., “landscape of fear”), which can further increase or even decrease local plant biomass (Schmitz et al. 2004).

On the other hand, little is known about the relative importance of direct and trait-mediated indirect effects of environmental stressors on population dynamics. Human activities are currently affecting various species negatively, and researchers tend to focus on the loss of biodiversity thorough the direct effects of environmental stressors on death or birth rates. But if some species change their traits in response to anthropogenic disturbances, it can alter the demography of other interacting species (i.e., the trait-mediated indirect effects) (Abrams 1995, Werner and Peacor 2003, Peacor et al. 2020). For example, human activities can invoke fear in some species and reduce their foraging behaviors (Smith et al. 2024), which can increase their prey’s performance (Leighton et al. 2010, Mehlhoop et al. 2022). While an increasing number of studies proposed the potential importance of trait-mediated indirect effects, it has in general been difficult to quantify the relative contributions of direct and indirect pathways of environmental stressor to focal species’ population dynamics.

Here we use a rotifer-alga experimental system (Fig. 1) for quantifying trait-mediated indirect effects. The rotifer-alga system has been used in various studies to understand species coexistence (Rothhaupt 1988), predator-prey population cycles (Fussmann et al. 2000, Verschoor et al. 2004, Scheuerl and Stelzer 2019, Blasius et al. 2020), and eco-evolutionary dynamics (Boraas et al. 1998, Fussmann et al. 2003, Yoshida et al. 2003, Yoshida et al. 2007, Becks et al. 2010, Becks et al. 2012, Hiltunen et al. 2014, Kasada et al. 2014, Declerck et al. 2015, Haafke et al. 2016, Hermann and Becks 2022). Particularly, many studies focused on clump formation as algal defense against rotifer predation (reviewed in Yamamichi 2020). In the freshwater unicellular green alga *Chlamydomonas reinhardtii*, the defense trait is palmelloid clump formation, which is too big to be eaten by gape-limited predators including rotifers (Lurling and Beekman 2006, Sathe and Durand 2016, Réveillon and Becks 2024), and such multicellularity is heritable and can evolve in laboratory experiments (Becks et al. 2010, Herron et al. 2019, Bernardes et al. 2021).

**Figure 1.**
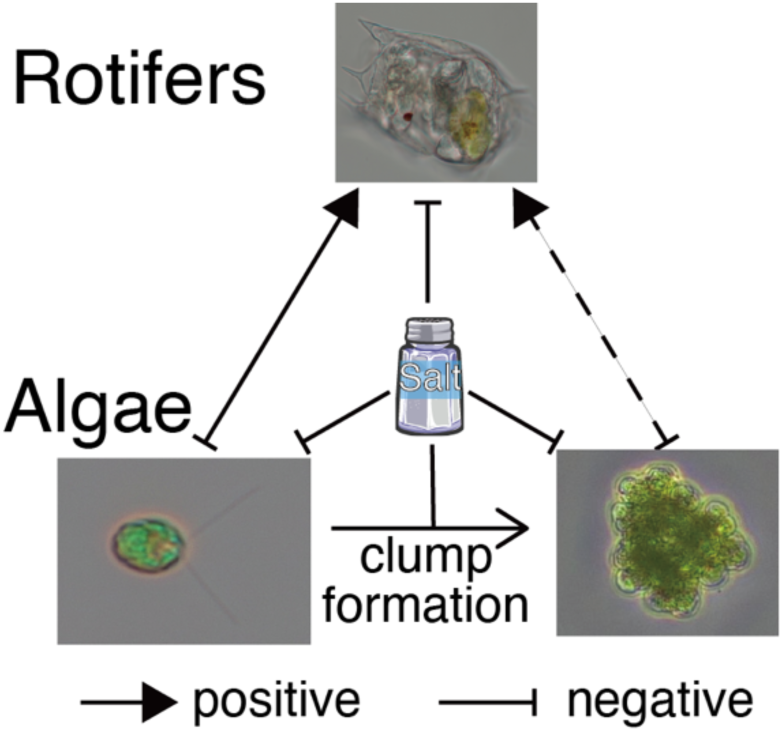
Schematic representation of the system. The green algae *Chlamydomonas sphaeroides* is consumed by rotifers *Brachionus calyciflorus* (the solid and dashed oblique arrows between them). While the salinity directly decreases the growth of both species (the three arrows from the salt), it also induces the clump formation of the green algae (the horizontal arrow). The clumps weaken the predator-prey interaction because rotifers are gape-limited predators (i.e., the trait-mediated indirect effect of salinity stress: the dashed oblique arrows). The salt icon by Servier https://smart.servier.com/ is licensed under CC-BY 3.0 Unported https://creativecommons.org/licenses/by/3.0/. Photos were taken by S. Shibasaki.

Importantly, rapid evolution driven by predation is not a sole cause of clump formation: clump formation in green algae is also induced by phenotypic plasticity triggered by environmental stressors (Cheloni and Slaveykova 2021, de Carpentier et al. 2022, Cornwallis et al. 2023) including salinity stress (Khona et al. 2016, Farkas et al. 2023). Human activities have salinized freshwater systems globally through, for example, roadway de-icing (Dugan et al. 2017, Kaushal et al. 2021, Cunillera-Montcusí et al. 2022), impeding population growth of freshwater phyto- and zooplankton species (Sarma et al. 2006, Venâncio et al. 2019). Therefore, human activities may not only directly increase the death rates of zooplankton species, but also indirectly harm them by inducing palmelloid clump formation of green algae.

Here, to understand trait-mediated indirect effects of salinity stress, we conducted laboratory experiments using four freshwater plankton species: a rotifer (*Brachionus calyciflorus*) as a predator and three green algae (*Chlamydomonas sphaeroides*, *Chlamydomonas reinhardtii*, and *Chlorella vulgaris*) as prey. *Chlamydomonas sphaeroides* is not a model organism, but phylogenetically close to *Chlamydomonas reinhardtii* (Yumoto et al. 2013) and was used as a food for *B. calyciflorus* in previous experiments (Rothhaupt 1988). We first show that *Chlamydomonas sphaeroides* formed remarkably larger clumps than the other two algal species under a moderate salinity stress (0.06M NaCl) by phenotypic plasticity. Then, we show that large clumps of *Chlamydomonas sphaeroides* were less likely to be consumed by rotifers by comparing the clump size distributions before and after exposure to predation under salinity stress. Finally, we show that the rotifer’s attack rate on the green alga became approximately half under salinity stress by Bayesian parameter estimation of the Lotka-Volterra model based on co-culture time-series data with and without salinity stress. These results indicate that salinity-induced clump formation can weaken predator-prey interactions in freshwater ecosystems, highlighting the importance of considering trait-mediated indirect effects of environmental stressors to understand community dynamics in changing environments.

## Materials and Methods

### Strain Information

The unicellular green algae, *Chlorella vulgaris* (NIES-227), *Chlamydomonas reinhardtii* (NIES-2236), and *Chlamydomonas sphaeroides* (NIES-2242), were obtained from the National Institute for Environmental Studies (NIES), Japan. The green algae were maintained in the C medium (Ichimura 1971) at 23℃ with the light cycle L:D = 14:10. The details of the culture conditions of the green algae were identical to our previous study (Shibasaki and Yamamichi 2024).

The facultative parthenogenetic rotifer *B. calyciflorus* (*elevetus*) (*sensu* Otake et al. 2025) was isolated from a water sample in Kanazawa Castle Park, Ishikawa prefecture, Japan, obtained in November 2023 by Dr. Yurie Otake at Kyoto University. The rotifer has been maintained in C medium with *Chlorella vulgaris* as prey under 25℃ with the light cycle 14:10. The rotifer culture was transferred into a fresh C medium with a weekly dilution rate ca. 10 % while adding 1 ml culture of *Chlorella vulgaris*.

Before adding the rotifers to experimental flasks, the stock culture of rotifers was filtered twice with a 40 µm cell strainer, left for rotifers to starve for four hours in the dark at 25℃, and then filtered and washed again so that *Chlorella vulgaris* did not contaminate the experimental flasks. This protocol follows a previous study with a similar experimental system (Hermann and Becks 2022).

### Comparing the clump sizes

We first compared the clump sizes of the three green algae in the following four conditions with three replicates in each: (i) control (C medium), (ii) with predators (adding rotifers at an initial density of ca. one female individual/ml), (iii) with salt (0.06M NaCl C medium), and (iv) with predators and salt (adding rotifers at initial density ca. one female individual/ml to 0.06M NaCl C medium). We grew the green algae (and rotifers, if applicable) in 50 ml of the media in a 300 ml Erlenmeyer flask for seven days at 25℃ with the light cycle L:D = 14:10 (using daylight fluorescent lamps with light intensity 145 ± 5 μmol m^−2^s^−1^) while shaking (60 rpm). After cultivation, we collected 1 ml of culture from each flask and added 10 μl of Lugol’s solution (20 g of potassium iodide and 10 g of iodine mixed in 200 ml of distilled water) to fix the green algae. We measured the sizes (i.e., the number of cells per clump) of 300 clumps per sample under microscopy.

### Evaluating salinity-induced clump formation as a defense trait

Next, we examined whether large clumps of *Chlamydomonas sphaeroides* induced by the salinity stress reduced rotifer predation. We first grew *Chlamydomonas sphaeroides* in 50 ml of 0.06M NaCl C medium for seven days in six 300 ml Erlenmeyer flasks so that the green algae formed clumps. Then, we sampled 5 ml of culture from each flask and fixed it using 50 μl of Lugol’s solution. We measured the sizes of 300 clumps in each. 5 ml of rotifer cultures (in 0.06M NaCl C medium) were added to three of the six flasks (i.e., test cases) so that the initial density of rotifers was 0.1 individual/ml, while we added 5 ml of fresh 0.06M NaCl C medium to the other three flasks (controls). The six cultures were grown for six hours under light (daylight fluorescent lamps with light intensity 145 ± 5 μmol m^−2^s^−1^) while shaking (60 rpm) at 25℃. We then sampled 5 ml of culture from each flask, fixed them with Lugol’s solution, and measured the clump sizes as explained above.

### Predator-prey dynamics and parameter estimation

To investigate how salinity stress alters predator-prey dynamics, we grew the rotifers and green algae (*Chlorella vulgaris* and *Chlamydomonas sphaeroides*) with and without salinity stress (C medium with 0.06M NaCl and 0M NaCl, respectively) for seven days. The plankton species were grown in 50 ml of the growth medium in a 300 ml-Erlenmeyer flask at 25℃ with the light cycle 14:10 (light intensity145 ± 5 μmol m^−2^s^−1^) while shaking (60 rpm). We estimated the population densities of green algae and rotifers daily by sampling 1 ml of culture from each flask. Because it was difficult to estimate the densities of *Chlamydomonas sphaeroides* with clumps using an automated cell counter Countess II FL (Thermo Fisher Scientific Inc.), we removed clumping as follows: we centrifuged the sampled cultures (4℃, 2120 x g, 10 min), removed the supernatant, added 1 ml of C medium to each, and left them for two hours at 25℃ under lights. This protocol resolved many clumps because the clump formation in this system was caused by phenotypic plasticity (Khona et al. 2016; see also Fig. S12). After two hours, we fixed the samples by adding 10 μl of Lugol’s solution to each. For each sample, we measured the densities of the green algae three times by Countess II FL (using 30 μl culture) and counted female rotifers under microscopy.

To distinguish the direct (i.e., increasing the mortality rate) and indirect (i.e., decreasing attack rate due to the salinity-induced clump formation of prey) impacts of salinity stress on the rotifer’s population dynamics, we fitted our experimental data to the following Lotka-Volterra predator-prey model with prey logistic growth and predator linear (Holling type I) functional response:

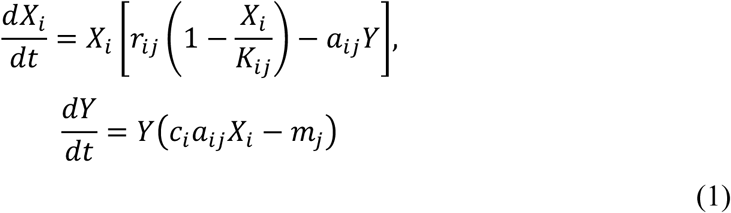

where 𝑋*_i_* is the scaled density of the green algae 𝑖 (cells/ml divided by 10^8^ for *Chlorella vulgaris* and 10^7^ for *Chlamydomonas sphaeroides*), 𝑌 is the scaled density of female rotifers (female individuals/ml divided by 10^3^), *r_ij_* is the intrinsic growth rate of the green algae *i* in the absence (*j* = 1) or presence (*j* = 2) of the 0.06M NaCl salinity stress, *K_ij_* is the carrying capacity of the green algae *i* in the absence (*j* = 1) or presence (*j* = 2) of the 0.06M NaCl salinity stress, *a_ij_* is the rotifer’s attack rate on green algae *i* in the absence (*j* = 1) or presence (*j* = 2) of the 0.06M NaCl salinity stress, *c_i_* is the conversion efficiency of the green algae *i* to the rotifer’s density, which we assume the salinity stress would not affect, and *m_j_* is the mortality rate of the rotifers in the absence (*j* = 1) or presence (*j* = 2) of the 0.06M NaCl salinity stress. We used the scaled densities of the prey and predator so that their maximum densities were about unity, which facilitated the following Bayesian parameter estimation.

Parameter values were estimated using Bayesian statistics (see Supporting Information for technical details). First, we estimated posterior distributions of the intrinsic growth rates (*r_ij_*) and the carrying capacities (*K_ij_*) of *Chlorella vulgaris* (*i* = 1) and *Chlamydomonas sphaeroides* (*i* = 2) in the absence (*j* = 1) or presence (*j* = 2) of the 0.06M NaCl salinity stress, respectively, from nine-day mono-culture experiments (Supporting Information S1.1-S1.2 and Figs. S1-S4). Next, we estimated the attack rate (*a*_1*j*_), conversion efficiency (*c*_1_), and mortality rate (*m_j_*) of rotifers in the absence (*j* = 1) or presence (*j* = 2) of the 0.06M NaCl salinity stress based on the seven-day co-culture experiments with *Chlorella vulgaris* (Supporting Information S1.3 and Figs. S5-S7). We assumed that the negative impact of salinity on rotifers came through the mortality rate only (i.e., *a*_11_ = *a*_12_) because *Chlorella vulgaris* seldom formed clumps in the presence of the 0.06M NaCl salinity stress (see Results). Finally, we estimated the attack rates on *Chlamydomonas sphaeroides* (*a*_2*j*_), conversion efficiency (*c*_2_), and mortality rate (*m_j_*) of rotifers in the absence (*j* = 1) or presence (*j* = 2) of the 0.06M NaCl salinity stress (Supporting Information S1.3 and Figs. S8-S10) utilizing the posterior distributions of *m_j_* inferred from the rotifer-*Chlorella* co-culture experiments as prior distributions.

### Experimental evolution of clumping formation

We also cultivated *Chlamydomonas sphaeroides* under salinity stress for 91 days to see whether larger clumps can evolve or not, because palmelloid clump formation might be adaptive under environmental stress (Tong et al. 2022). Previous studies showed that palmelloid clump formation of *Chlamydomonas reinhardtii* can evolve rapidly in laboratory experiments (Becks et al. 2010). Although clump formation of *Chlamydomonas sphaeroides* is phenotypic plasticity (see Results), we expected that we might be able to observe genetic assimilation, where clumps originally formed in response to salinity stress later become genetically fixed via selection (Pigliucci et al. 2006). Such evolved strains would allow us to investigate whether larger clumps were more unlikely to be consumed by rotifers or not in the absence of salinity stress. We grew the ten lines of the green algae in 20 ml of C medium in 200 ml Erlenmeyer flasks at 25℃ with the light cycle 14:10 (light intensity 145 ± 5 μmol m^−2^s^−1^) while shaking (120 rpm). Within the ten lines, five lines were grown under salinity stress (0.06M NaCl) while the rest five were without salt (0M NaCl). The initial algal density was approximately 1×10^5^ cells/ml. Every week, we collected 200 μl of culture medium from each flask, centrifuged the sampled cultures (4℃, 2120 x g, 10 min), removed the supernatant, and added each sample to 20 ml of fresh C medium with or without NaCl. Note that we accidentally sampled 1000 μl, instead of 200 μl, of cultures and transferred into fresh media on day 21.

We weekly measured algae densities and mean diameter of particles (i.e., either single cells or multi-cellular clumps) using an automatic cell counter analyzer, CASY TT (Innovatis). In addition, we also examined whether *Chlamydomonas sphaeroides* maintained palmelloid clump formation when salinity stress was removed as follows: we collected 5 ml of culture medium from each line under salinity stress, centrifuged the sampled cultures, removed the supernatant, and grew in 2 ml of fresh C medium with and without NaCl, respectively, for six hours. After six hours, we measured the mean diameter of the particles by CASY TT. If the green algae lost phenotypic plasticity, the mean diameter without salinity stress was expected to be as high as with salinity stress. In contrast, the six-hour culture under salinity stress would not affect the mean diameters. Note that a part of the data on the mean diameter is missing on days 42 and 91, because we could not use CASY TT due to its malfunctioning on December 31st, 2024, and February 18th, 2025, respectively.

### Statistical analyses

All statistical analyses were performed in R version 4.3.1 (R Core Team 2024). The effects of salt and rotifers on clump sizes of the green algae were analyzed by the non-parametric two-way analysis of variance (ANOVA) with the aligned rank transformation in ARTool library version 0.11.1 (Kay et al. 2025). The linear mixed model was implemented by lmerTest library version 3.1.3 (Kuznetsova et al. 2017). The parameter values in the predator-prey model were estimated by RStan library version 2.32.6 (Stan Development Team 2025). In the main text, we reported the mean values and 95% credible intervals (CI) of the posterior distributions of the fold changes in the attack rate on *Chlamydomonas sphaeroides* and the rotifer mortality rate.

## Results

*Chlamydomonas sphaeroides* generates large clumps under salt stress We first compared the clump sizes of the three freshwater green algae, *Chlorella vulgaris*, *Chlamydomonas reinhardtii*, and *Chlamydomonas sphaeroides*, in the presence of either or both of two stressors: salt and predators (rotifers). Although all four taxa in this study are freshwater species, they can survive in the C medium containing 0.06M NaCl, which is about 10% seawater salinity (see Figs. S1-S10 as examples) (Shibasaki and Yamamichi 2024). Figure 2 shows that the median clump sizes of the three green algae were ∼1.0 in the control cases, confirming that they were unicellular algae. While *Chlorella vulgaris* remained essentially unicellular in the presence of salt and rotifers, the two *Chlamydomonas* species increased clump size in the presence of salt. The median clump size of *Chlamydomonas reinhardtii* increased from one to two cells due to the salinity stress, but their typical clump sizes were less than four cells per clump. *Chlamydomonas sphaeroides* increased in median clump size from one to eight or more cells under salinity stress, and we even observed more than 30 cells in a single clump (see the bottom panels of Fig. 2). Statistical analyses also suggest that salinity had a large impact on the clump sizes of *Chlamydomonas sphaeroides* (Table 1). Because the green algae immediately became unicellular again when the salinity stress was removed (Fig. S12), the palmelloid clump formation in response to salinity stress seems to be phenotypic plasticity, not evolution.

**Figure 2.**
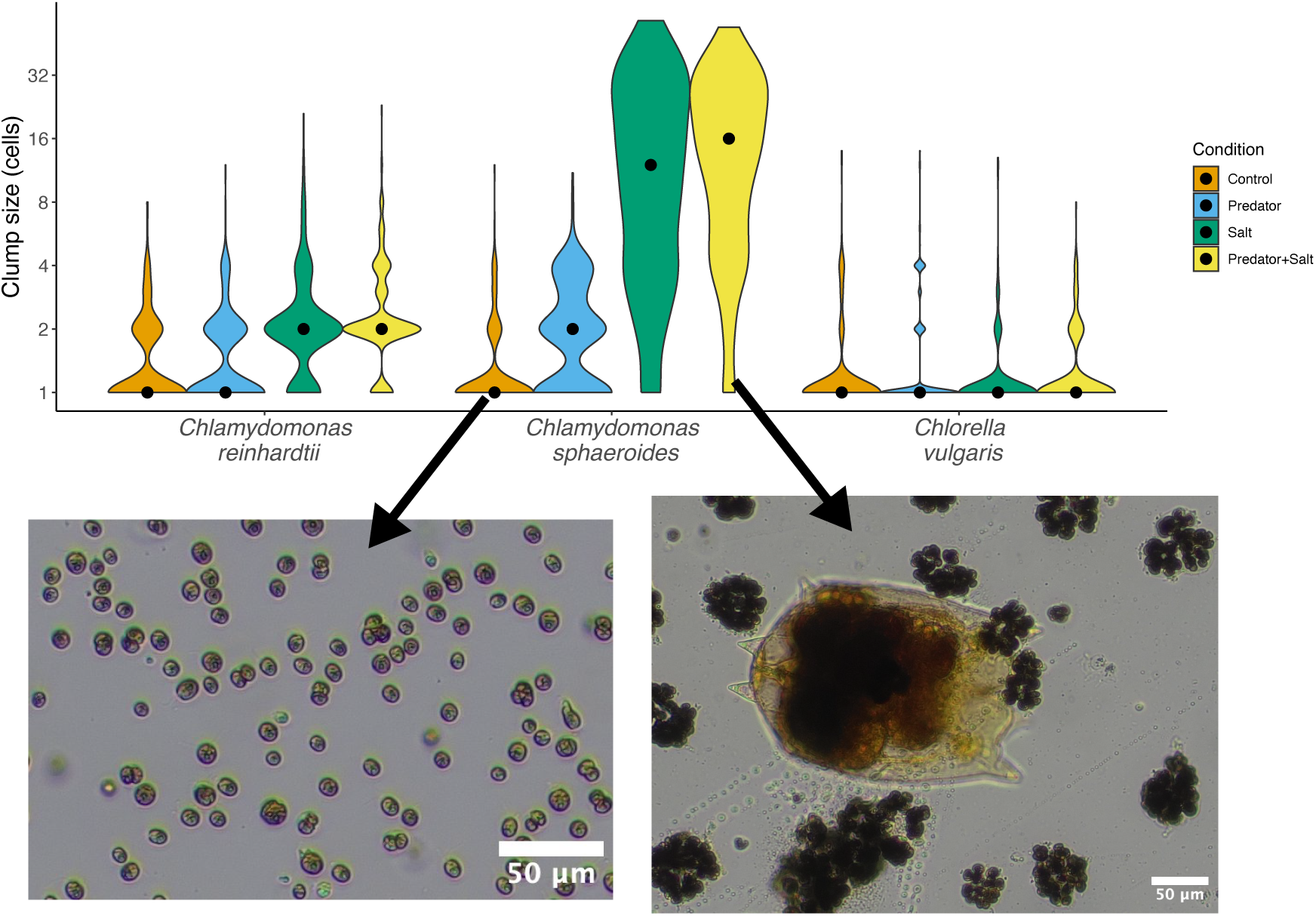
Large clump formation was induced by salinity stress in *Chlamydomonas sphaeroides*. Top: The distributions of clump sizes of *Chlamydomonas reinhardtii* (left), *Chlamydomonas sphaeroides* (middle), and *Chlorella vulgaris* (right) under four conditions (orange: control, blue: with rotifers, green: with 0.06M NaCl, and yellow: with rotifers and 0.06M NaCl) after seven days. The black dots represent the median clump sizes of 900 clumps. Bottom: examples of clump sizes of *Chlamydomonas sphaeroides* in the control condition (left) and in the presence of salinity stress and rotifers (right). The white bars represent the scale (50 μm). Photos were taken by S. Shibasaki.

**Table 1.** Non-parametric two-way ANOVA and its effect size (partial *η^2^*). Df is the degree of freedom.

| Species | Stressor | Df | Df.res | *F* value | *p* value | partial $\eta^2$ |
| --- | --- | --- | --- | --- | --- | --- |
| *Chlamydomonas sphaeroides* | Salt | 1 | 1196 | 1626.542 | 0 | 0.576 |
|  | Predator | 1 | 1196 | 65.765 | 0 | 0.052 |
| | Salt ×<br>Predator | 1 | 1196 | 21.373 | 0 | 0.018 |
| *Chlamydomonas reinhardtii* | Salt | 1 | 1196 | 241.7099 | 0 | 0.168 |
|  | Predator | 1 | 1196 | 14.1246 | 0 | 0.012 |
| | Salt ×<br>Predator | 1 | 1196 | 3.3138 | 0.069 | 0.003 |
| *Chlorella vulgaris* | Salt | 1 | 1196 | 7.1755 | 0.008 | 0.006 |
|  | Predator | 1 | 1196 | 20.3152 | 0 | 0.017 |
| | Salt ×<br>Predator | 1 | 1196 | 4.5742 | 0.033 | 0.004 |

### Larger clumps impeded the predation by the rotifers

Next, we examined whether large clumps of *Chlamydomonas sphaeroides* under salinity stress could work as a defense trait against gape-limited predation by rotifers. We addressed this question by comparing the clump size distribution before and after exposure to rotifer predation for six hours. Figure 3 shows how the distributions of clump sizes changed with (Fig. 3A) and without (Fig. 3B) rotifers. In the absence of rotifers, the median clump sizes decreased or were little changed in six hours (Fig. 3B). On the other hand, the median clump sizes clearly increased in the presence of rotifers (Fig. 3A). The linear mixed model suggested that the clump sizes, on average, decreased in six hours but increased if the green algae co-cultured with the rotifers (Table 2). The violin plots show that the proportion of small clumps (four or fewer cells per clump) decreased with predators. Because the doubling time of *Chlamydomonas sphaeroides* under salinity was estimated to be log2/*r*_22_ ≈ 1.33 days (see Figs. S3-S4), which is longer than six hours, the presence of rotifers would not induce the formation of larger palmelloid clump in this experiment. Thus, the rotifer predation would decrease the proportion of small clumps, but not large ones.

**Figure 3.**
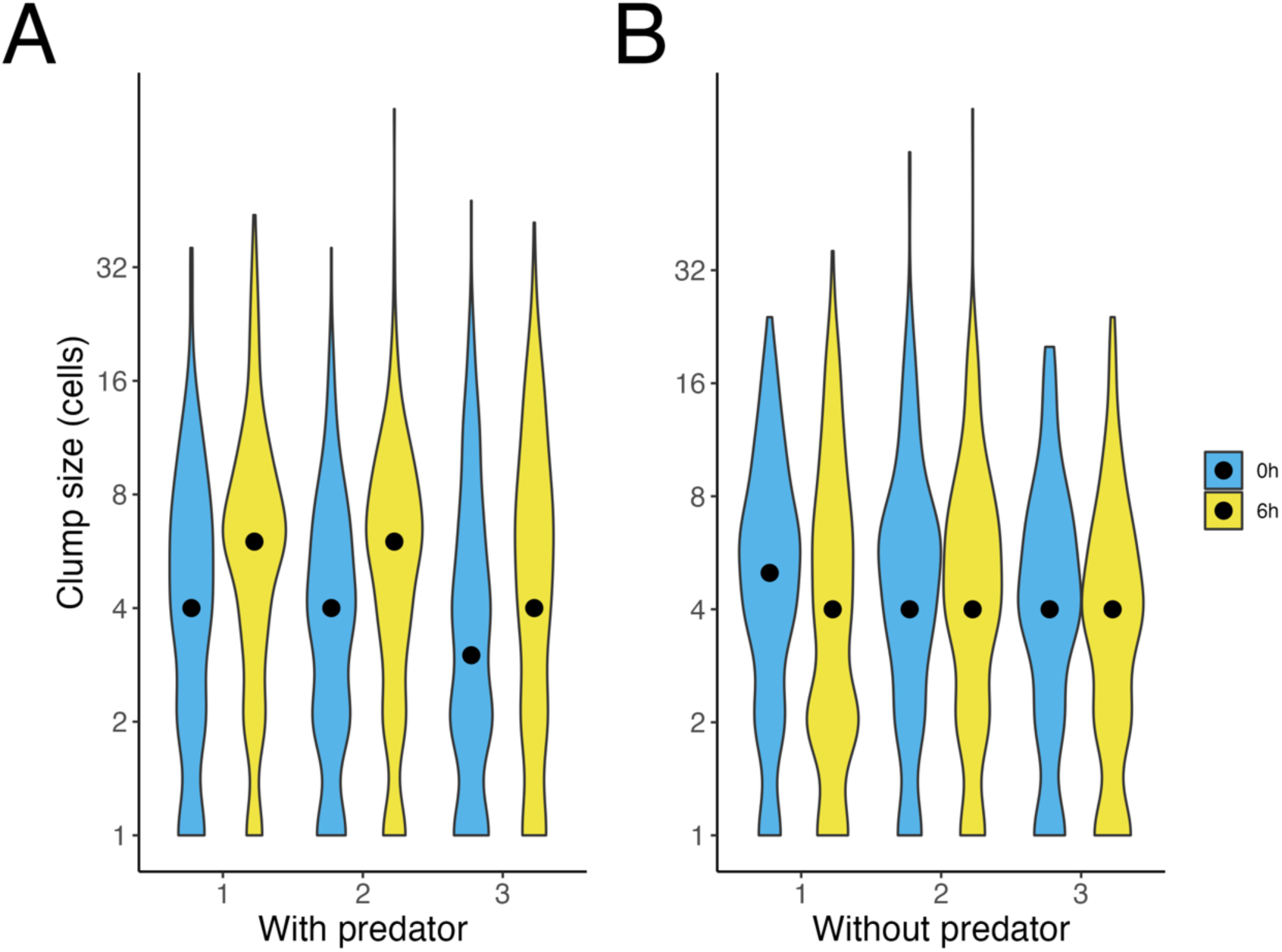
Large clumps were not consumed by rotifers. The distributions of clump sizes of *Chlamydomonas sphaeroides* in the presence (A) or absence (B) of rotifers in 0.06M NaCl C medium with three replicates (flasks). By comparing the size distributions before (blue) and after (yellow) the exposure to rotifers for six hours, it is clear that rotifers consumed smaller clumps (A) and salinity stress cannot explain the difference (B). The black dots represent the median clump size of 300 clumps.

**Table 2.** Fixed effect in the linear mixed model on the clump sizes after predation. S.E. is standard error and Df is the degree of freedom.

| Variable | Explanation | Estimate | S.E. | Df | *t* value | *p* value |
| --- | --- | --- | --- | --- | --- | --- |
| Intercept |  | 2.054 | 0.081 | 5.157 | 25.307 | 0 |
| Time | Six hours passed | −0.169 | 0.056 | 3592 | −3.015 | 0.003 |
| Predators | Presence of rotifers | −0.27 | 0.115 | 5.157 | −2.354 | 0.064 |
| Time ×<br>predators |  | 0.343 | 0.115 | 5.157 | 2.991 | 0.029 |

We then examined how salinity stress changed predator-prey dynamics by conducting co-culture experiments of the green algae and rotifers for seven days. While the population dynamics of the green algae *Chlamydomonas sphaeroides* did not seemingly change as a function of salinity stress (Fig. 4A), we observed a remarkable decrease in rotifer’s population growth in the presence of salinity stress (Fig. 4B). To disentangle the relative importance of direct and trait-mediated indirect impacts of salinity stress, we fitted the experimental data to the Lotka-Volterra predator-prey model (Eq. 1) by performing the Bayesian estimation (see Supporting Information and Figs. S1-S10 for more details). Fig. 4C shows that the mortality rate of the rotifers under salinity stress (*m*_2_) was more than 17 times (95% CI [6.07, 58.27]) larger than that without salinity stress (*m*_1_). The attack rate on *Chlamydomonas sphaeroides* under salinity stress (*a*_22_) was, on the other hand, about half (95% CI [0.18, 0.98]) of that in the absence of salinity stress (*a*_21_). Owing to the low rotifer density, the posterior distribution of *a*_22_ remained close to its prior and exhibited substantial uncertainty, whereas the posterior distribution of *a*_21_ shifted markedly from its prior (Fig. S10). These results indicate that the attack rates on *Chlamydomonas sphaeroides* are affected by salinity. Taken together, we successfully quantified the direct and trait-mediated indirect impacts of salinity stress on the rotifer population dynamics, and the direct impact was more severe.

**Figure 4.**
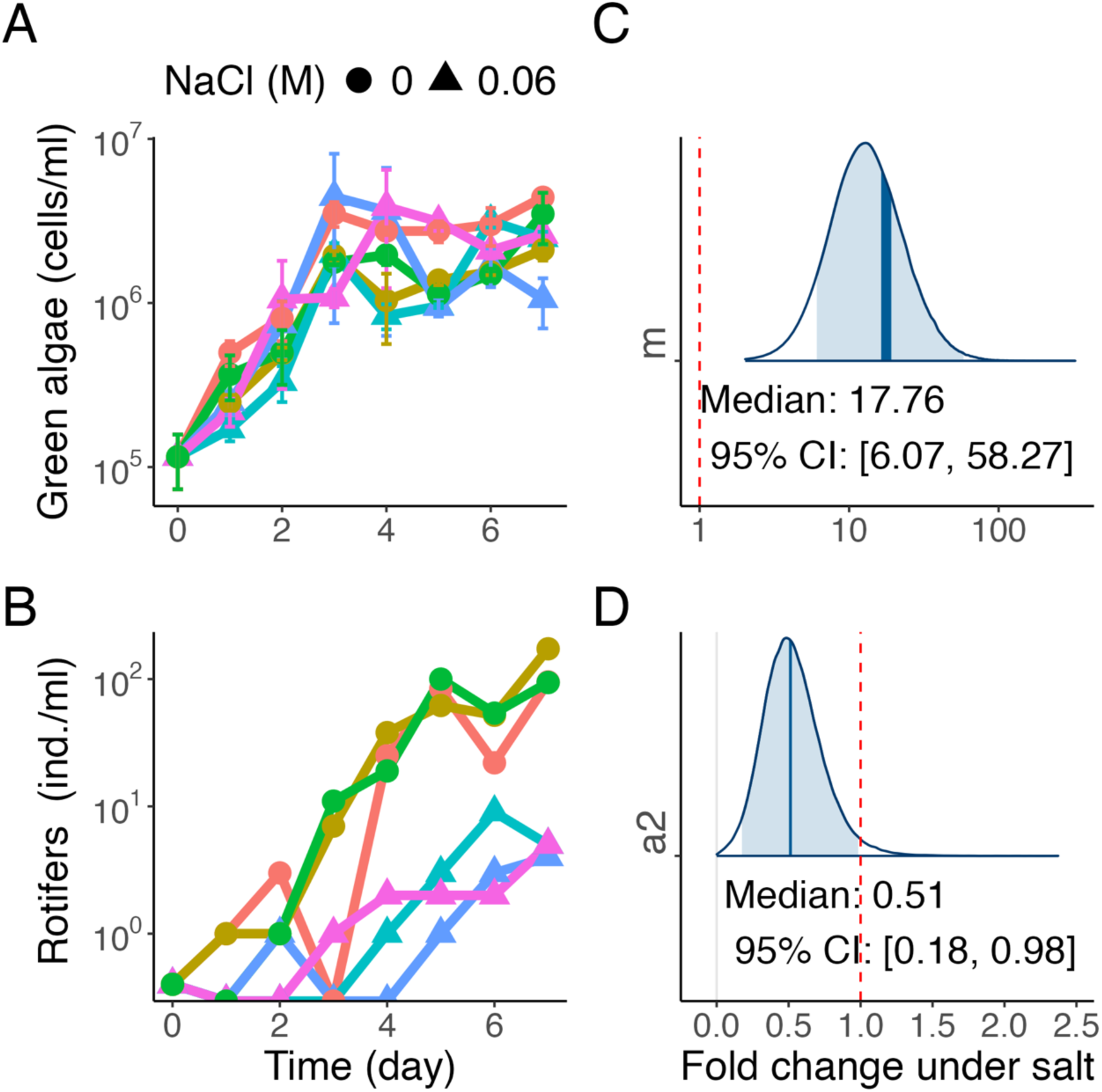
Predator-prey dynamics with or without salinity stress. A and B: Population dynamics of *Chlamydomonas sphaeroides* (A) and the rotifers (B) are shown. The shapes of markers represent the salinity stress (circle: 0M NaCl, and triangle: 0.06M NaCl). Under each salinity stress, we performed a one-week experiment with three replicates (shown by different colors). The error bars in panel A represent the standard errors, while the markers show the mean values. C and D: The posterior probability distributions of the fold changes of the rotifer’s mortality rate (*m*_2_/*m*_1_: C) and attack rates on *Chlamydomonas sphaeroides* (*a*_22_/*a*_21_: D) are shown, respectively. The vertical blue lines show the median values, and the shaded areas show 95% CIs. The vertical red dashed lines show one (i.e., no fold changes).

### The salinity stress did not lead to the evolution of larger clumps or loss of phenotypic plasticity

We also examined whether the clump size evolved or not when *Chlamydomonas sphaeroides* were grown under salinity for 91 days. Fig. S11 shows that, while all five lines without salinity stress (i.e., controls; orange in Fig. S11A) grew constantly during the experiment, the lines under salinity stress (blue in Fig. S11A) gradually decreased in growth rate from day 7 with some variation: one line (Salt-1) maintained a moderate population density (approximately 1-2×10^5^ cells/ml) during the experiment, whereas another (Salt-4) continued to decrease its density from day 70 and went extinct (its density was <100 cells/ml) at the end of the experiment. Fig. S11B shows the dynamics of mean diameters of the particles (single cells and multicellular clumps) during the experiment. Without salinity stress, the mean diameter remained around 10 μm, suggesting that the green algae were unicellular. On the other hand, the mean diameter under salinity stress was about 20 μm. This indicates that, if a single cell size did not change, a single clump would be composed of 2^3^ = 8 cells on average, which is consistent with our observation under microscopy (Figs. 1-2). However, there was no evidence that clump size evolved to be larger under salinity stress.

We also investigated whether *Chlamydomonas sphaeroides* maintained clump formation without salt during the experiment by weekly transferring the green algae into fresh C media with and without NaCl for six hours. Fig. S12 shows the mean diameter returned to approximately 10 μm without salinity stress, while the mean diameter remained 20 μm with the 0.06M NaCl salinity stress. These results indicate that the green algae returned to single cells when the salinity stress was removed, and that phenotypic plasticity was maintained during the 91-day experiment.

## Discussion

While human activities can directly threaten organisms by harvesting or destroying their habitats, these impacts can also propagate through communities by altering species interactions: i.e., trait-mediated indirect effects (Abrams 1995, Werner and Peacor 2003). Despite its potential importance, the relative impacts of direct and indirect effects on population dynamics were unclear. Here, we used a rotifer-alga system to quantify a trait-mediated indirect effect in freshwater communities. We found that the 0.06M NaCl salinity stress induced the algal clump formation (Fig. 2), and it weakened the interspecific interaction (predation) between the green algae and rotifers (Figs. 3-4). In freshwater systems, salinization is a severe problem caused by human activities (Dugan et al. 2017, Kaushal et al. 2021, Cunillera-Montcusí et al. 2022). Our results demonstrate the importance of a quantitative understanding of how environmental stressors can affect community dynamics via trait modifications.

Previous studies investigated the clump formation of *Chlamydomonas reinhardtii* as a defense trait against predation (Lurling and Beekman 2006, Becks et al. 2010, Becks et al. 2012, Sathe and Durand 2016, Bernardes et al. 2021). However, green algae form multicellular clumps under other kinds of stress (Cheloni and Slaveykova 2021, de Carpentier et al. 2022, Cornwallis et al. 2023) including salinity stress (Khona et al. 2016, Farkas et al. 2023). Our study shows that *Chlamydomonas sphaeroides* was sensitive to the salinity stress and formed larger clumps than two other model green algae, *Chlorella vulgaris* and *Chlamydomonas reinhardtii*, under 0.06M NaCl (Fig. 2). Salinity-induced clump formation was caused by phenotypic plasticity because the green algae immediately lost the clumps and became unicellular when the salinity was removed from the media (Fig. S12). The green algae formed clumps without predators (Figs. 2 and S11B), and thus their effect on the predator-prey interaction seems a by-product of a response to salinity. Although Tong et al. (2022) argued that the salinity-induced clump formation can be adaptive under stressful environments, our evolution experiment did not support this claim. Nevertheless, the salinity-induced clump formation worked as a defense trait against predation by rotifers as shown in previous studies (Becks et al. 2010, Réveillon and Becks 2024): our experiments showed that the rotifers were unlikely to consume large clumps (Fig. 3) and their attack rate on *Chlamydomonas sphaeroides* decreased under salinity (Fig. 4D). Our results, therefore, indicate that salinity stress weakened the predator-prey interaction by inducing the algae clump formation.

We found that the trait-mediated indirect effect of salinity stress (decreasing the attack rate on *Chlamydomonas sphaeroides a*_2*j*_ by half) was less intense than the direct effect (increasing the rotifer’s mortality rate *m_j_* by 17-fold) (Fig. 4). However, when we extrapolate the results, we found that it can have a significant effect: in the Lotka-Volterra predator-prey model (Eq. 1), the predators cannot persist if *c_i_a_ij_K_ij_* < *m_j_*. Hence, salinity stress can promote rotifer extinction by reducing *a_ij_* through algal clump formation as well as increasing *m_j_* and decreasing *K_ij_*. This is in stark contrast to the idea of indirect evolutionary rescue, where prey defense evolution prevents predator extinction (Yamamichi and Miner 2015, Cortez and Yamamichi 2019, Hermann and Becks 2022). Hermann and Becks (2022) showed that increasing salinity reduces the rotifer density through the direct effect, and the reduced predation pressure selects for less defended *Chlamydomonas reinhardtii* due to the growth-defense trade-off (Becks et al. 2010, Réveillon and Becks 2024). Because of the reduced frequency of defended algae, in turn, rotifer populations may be rescued from extinction. In our system, clump formation is induced by salinity stress, suggesting that rotifers may not be rescued from extinction. It will be interesting to integrate these two opposite consequences of increasing salinity for better understanding how clump formation under salinity stress can pleiotropically affect salt tolerance and predator-prey dynamics.

Expanding our experimental system may provide deeper insights into how human salinization activities disturb freshwater communities. For example, we will be able to combine two or more stressors (e.g., pH, temperature) to experimentally show how realistic environmental changes alter ecological dynamics (Kawaguchi and Yamamichi 2025). Although we focused on interactions between two species, we can introduce another green alga that may be less sensitive to salinity stress (e.g., *Chlorella vulgaris*) to the rotifer-*Chlamydomonas* system to investigate how salt changes the outcome of exploitative and apparent competition between the two algae species. Another possibility is introducing a zooplankton species that consumes (e.g., *Asplanchna brightwelli*) (Verschoor et al. 2004) or competitor of *B. calyciflorus* (e.g., *B. rubens*) (Rothhaupt 1988). Such expansion would allow us to investigate how salinization would change the stability and structure of complex food webs. Shibasaki and Terui (2024) recently showed that species richness non-linearly alters food chain length. Because salinity stress would reduce the number of freshwater predator species by increasing mortality rates and causing the trait-mediated indirect effects, it may be possible to observe the nonlinear relationship between species richness and food chain length by manipulating salinity in our freshwater systems in future studies.

In conclusion, the present study demonstrates that the rotifer-alga system is useful for investigating trait-mediated interaction effects caused by environmental stressors. We focused on the salinization of freshwater systems and analyzed how salinity stress changes the predator-prey interaction between green algae *Chlamydomonas sphaeroides* and rotifers *B. calyciflorus*. In addition to the direct effect of salinity stress increasing rotifer mortality rate, salinity stress induced the multicellular clump formation of the green algae, which worked as a by-product defense trait against gape-limited predation by rotifers. Therefore, the salinity impeded the growth of rotifers both directly and indirectly. Future studies will be able to investigate how freshwater salinization alters food webs by extending this experimental system.

## Supporting information

Supporitng Information

## Acknowledgment

The three strains of the green algae *Chlorella vulgaris* (NIES-227), *Chlamydomonas reinhardtii* (NIES-2236), and *Chlamydomonas sphaeroides* (NIES-2242), were obtained from the National Institute for Environmental Studies (NIES), Japan. We thank Yurie Otake for sharing the water sample from which we isolated *Brachionus calyciflorus*. We also appreciate Mayumi Aono for helping with the experiments as a technician. This study was supported by the Japan Society for the Promotion of Science (JSPS) Grant-in-Aid for Scientific Research (KAKENHI) JP22H02688, JP22H04983, and JP25K02340, Japan Science and Technology Agency (JST) CREST JPMJCR23N5, Inamori Research Grant 2024, and Research Organization for Information and Systems (ROIS) Research Grant of Strategic Research Project 2024-SRP-10 to M. Y.

## Author contribution

S.S. and M. Y. conceived the project, S.S. performed the experiments, conducted the statistical analyses, and wrote the initial draft. M.Y. supervised the project and acquired the funding. Both authors reviewed and edited the manuscript.

## Data availability

The experimental codes and programming codes are available from the following Zenodo repository: https://doi.org/10.5281/zenodo.17875555.

