## Supplementary material for "Quantifying a trait-mediated indirect effect of an environmental stressor on predator-prey dynamics": Supporitng Information

### S1: Parameter estimation

In this section, we explain the details of the procedure for estimating the parameter values in the Lotka-Volterra predator-prey model (Eq. 1) from the experimental data of rotifer-alga dynamics. We first assume that observation errors are log-normally distributed (S1.1), and estimate algal parameters (the intrinsic growth rate and carrying capacity) based on mono-culture experiments of *Chlorella vulgaris* and *Chlamydomonas sphaeroides* (S1.2, Figs. S1-S4). Then, we estimate rotifer parameters (the attack rate, conversion efficiency, and mortality rate) based on co-culture experiments (S1.3, Figs. S5-S10).

#### S1.1: Estimation of the population densities

We assume that the observed predator and prey densities are noisy measurements of the true densities. Specifically, observed rotifer and alga densities were modeled as log-normally distributed around the true population densities. This formulation captures the idea that deviations are proportional to the true values, i.e., a constant fold-change error, which is often more realistic for ecological count data than assuming constant additive noise. Mathematically,

$$x_i \sim \mathrm{LogNormal}(\log X_i, \sigma_1^2)$$
$$y \sim \mathrm{LogNormal}(\log Y, \sigma_2^2),$$

where $x_i$ and $y$ are observed scaled densities of the green algae $i$ (cells/ml divided by $10^8$ for *Chlorella vulgaris*, $i = 1$, and $10^7$ for *Chlamydomonas sphaeroides*, $i = 2$) and rotifers (female individuals/ml divided by $10^3$), respectively, $X_i$ and $Y$ are true scaled ones, and $\sigma_i$ reflects the observation errors ($i = 1, 2$). We assigned a half-normal prior with a scale of 0.5 to $\sigma_i$. This prior distribution corresponds to multiplicative observation errors such that the observed values were expected to lie between roughly 0.37 and 2.72 times the true population densities (covering approximately the 95% credible interval).

### S1.2: Estimation of the alga parameters

We estimated the intrinsic growth rates $r_{ij}$ and the carrying capacities $K_{ij}$ of *Chlorella vulgaris* ($i$ = 1) and *Chlamydomonas sphaeroides* ($i$ = 2), respectively, in the absence ($j$ = 1) or presence ($j$ = 2) of salinity stress from nine-day mono-culture experiments. We performed the mono-culture experiment with three replicates for each condition. We chose the following Cauchy and normal distributions as prior distributions of the intrinsic growth rates and the carrying capacities, respectively:

$$r_{ij} \sim \text{Cauchy}\ (0, 2.5)$$
$$K_{ij} \sim N\ (1, 0.2^2)$$

The hyperparameters of the prior distribution of $r_{ij}$ were set so that the prior distributions were broad and non-informative. The prior distribution of $K_{ij}$ corresponds to the fact that we scaled the algal densities so that their maximum scaled densities were around one. We ran 50,000 Markov Chain Monte Carlo (MCMC) iterations for each estimation with 5,000 warm-ups and four chains.

The posterior distributions of the parameter values and the fitting results are shown in Figs. S1-S4. In the following estimation, we fixed the values of $r_{ij}$ and $K_{ij}$ as the median values of the posterior distributions in the mono-culture, namely, *Chlorella* without salt: $r_{11}$ = 1.24 and $K_{11}$ = 0.43, *Chlorella* with salt: $r_{12}$ = 1.10 and $K_{12}$ = 0.64 (Figs. S1-S2), *Chlamydomonas* without salt: $r_{21}$ = 0.66 and $K_{21}$ = 0.62, and *Chlamydomonas* with salt: $r_{22}$ = 0.52 and $K_{22}$ = 0.43 (Figs. S3-S4).

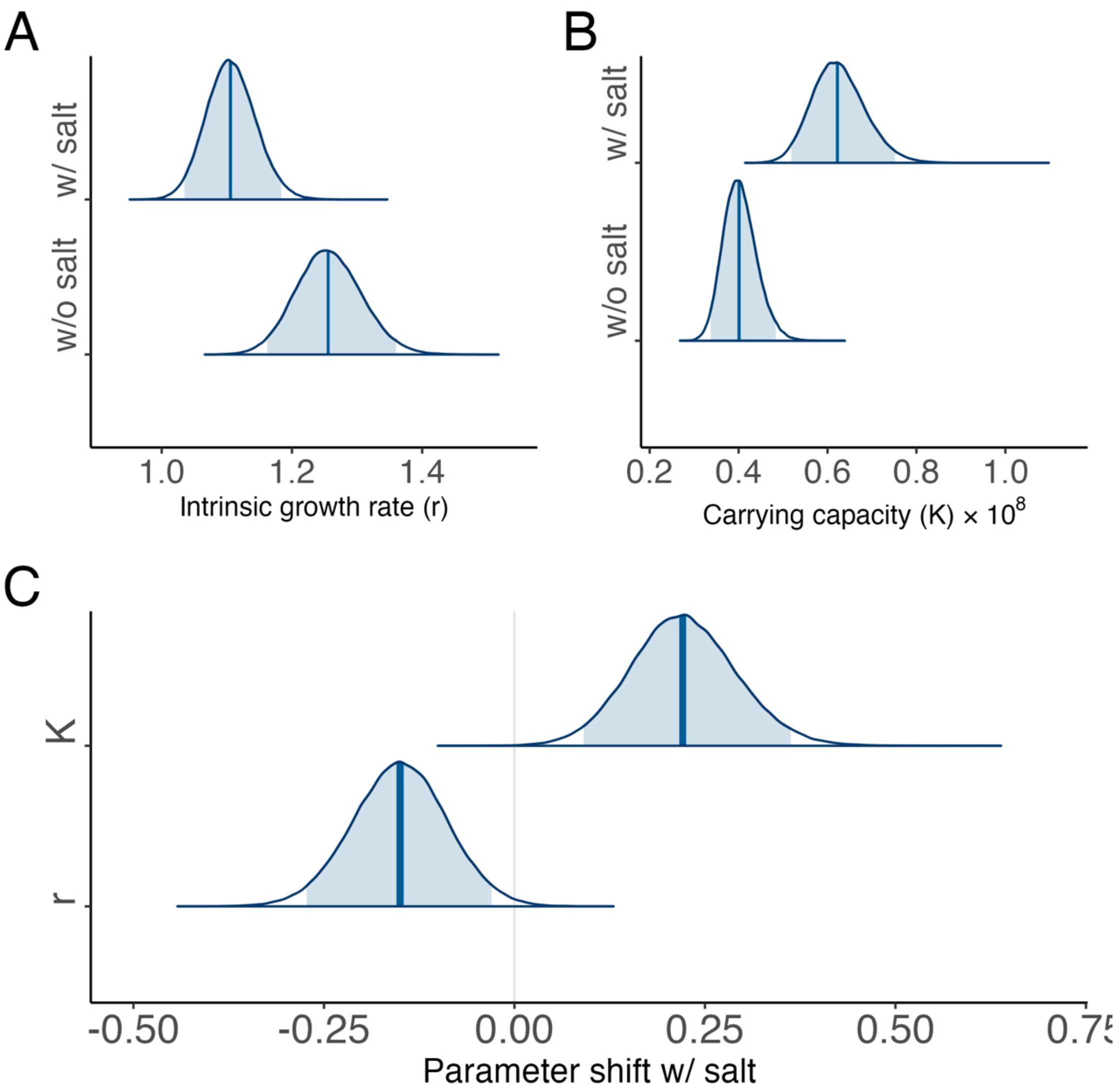


**Figure S1: Posterior distributions of the parameter values of *Chlorella vulgaris*.** A and B: posterior distributions of $r_{1j}$ (A) and $K_{1j}$ (B), respectively, where $j$ indicates the absence ($j$ = 1) or presence ($j$ = 2) of salinity stress. C: The posterior distributions of the $r_{12} – r_{11}$ and $K_{12} – K_{11}$, respectively. Salinity stress decreased the intrinsic growth rate but increased the carrying capacity in *Chlorella vulgaris*. In all panels, the vertical lines show the median values, while the shaded areas represent the 95% credible intervals (CI).

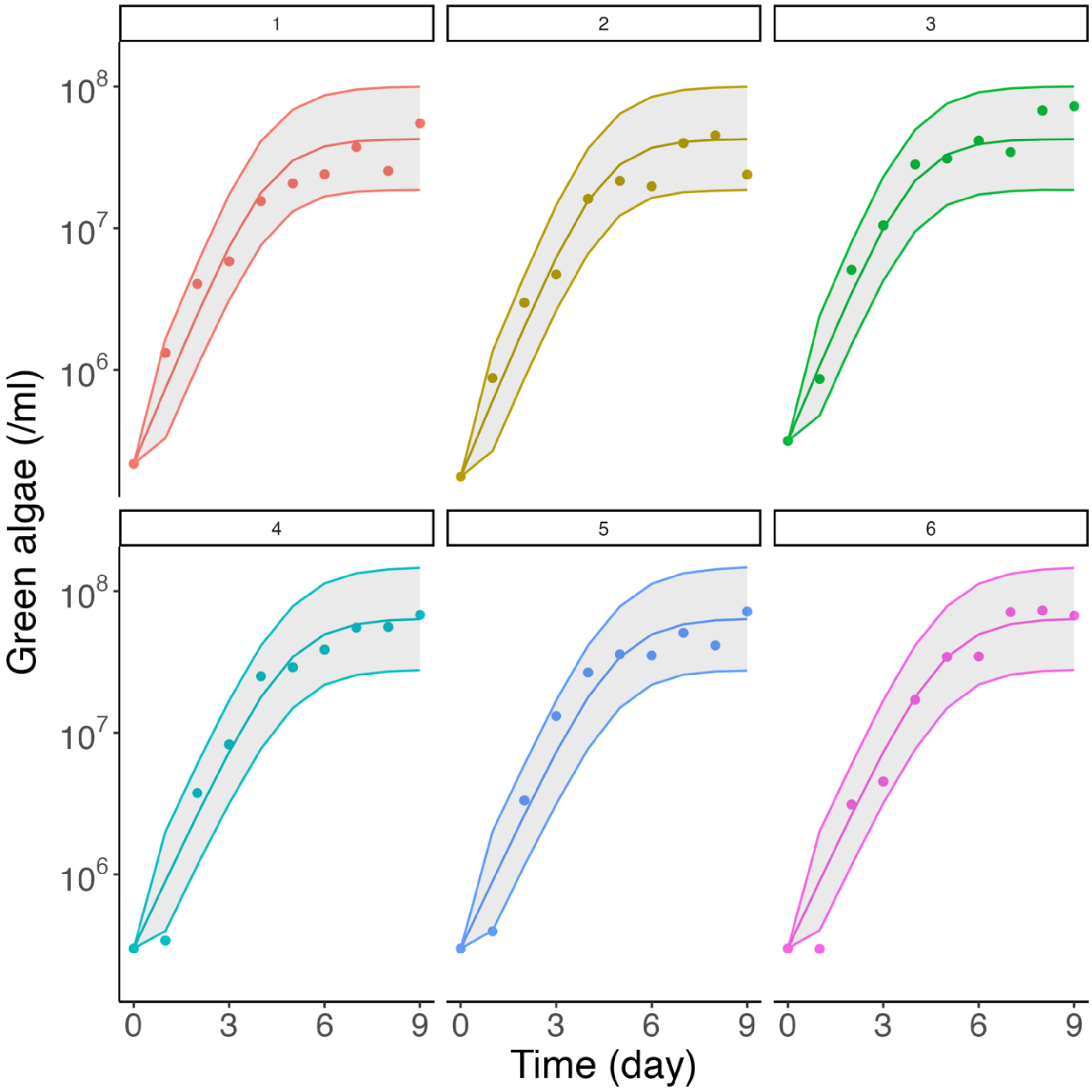


**Figure S2: Population dynamics of *Chlorella vulgaris* in mono-culture.**
The population dynamics of *Chlorella vulgaris* with (the top three panels) and without (the bottom three panels) salinity stress, with three replicates in each condition. The dots represent the mean densities of the green algae from three measurements, the shaded areas show the 95% CI, and the middle-colored lines indicate the dynamics with the median values from the posterior distribution.

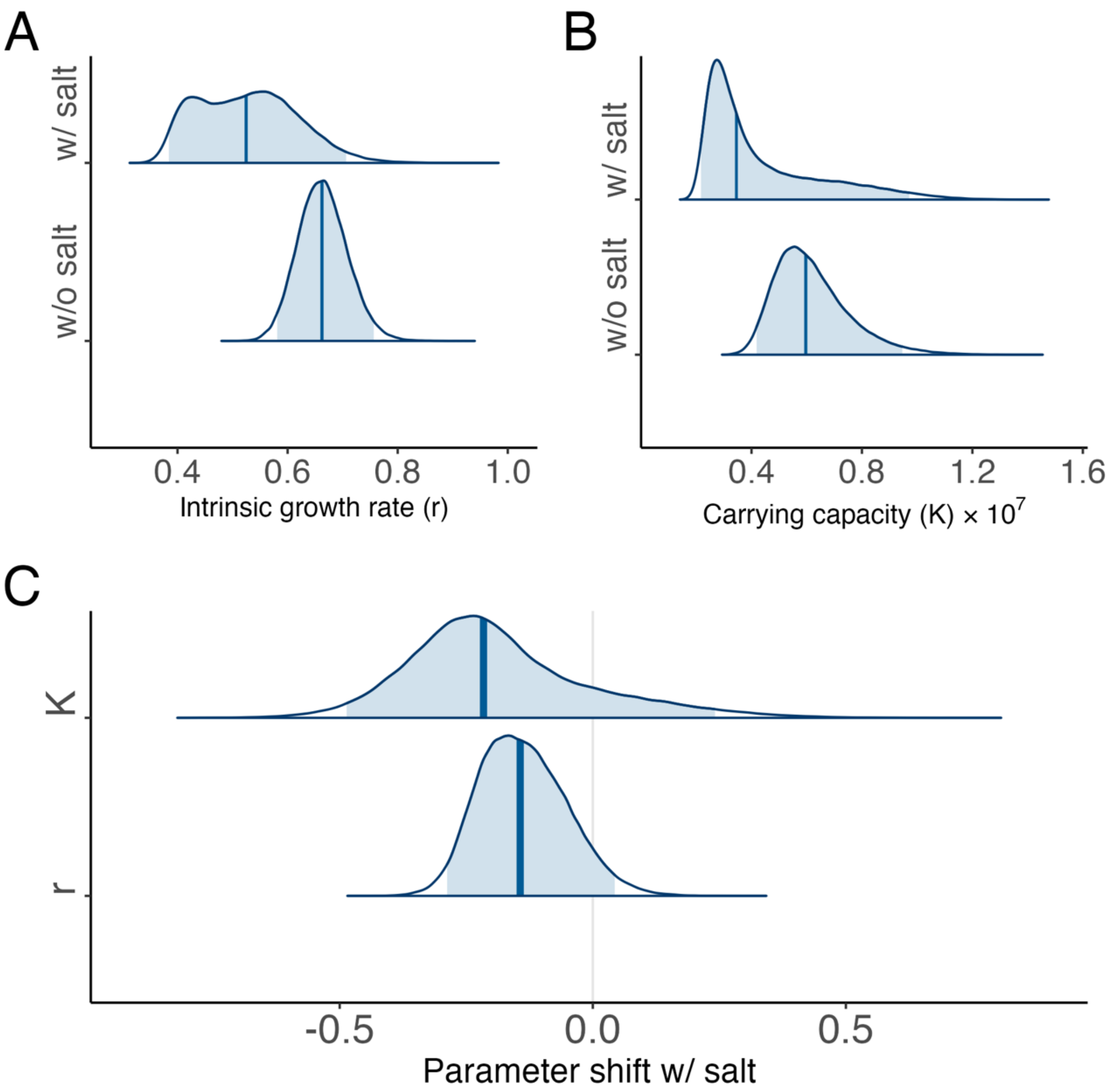


**Figure S3: Posterior distributions of the parameter values of *Chlamydomonas sphaeroides*.** A and B: posterior distributions of $r_{2j}$ (A) and $K_{2j}$ (B), respectively, where $j$ indicates the absence ($j$ = 1) or presence ($j$ = 2) of salinity stress. C: The posterior distributions of the $r_{22} - r_{21}$ and $K_{22} - K_{21}$, respectively. Salinity stress decreased both the intrinsic growth rate and carrying capacity in *Chlamydomonas sphaeroides*. In all panels, the vertical lines show the median values, while the shaded areas represent the 95% CI.

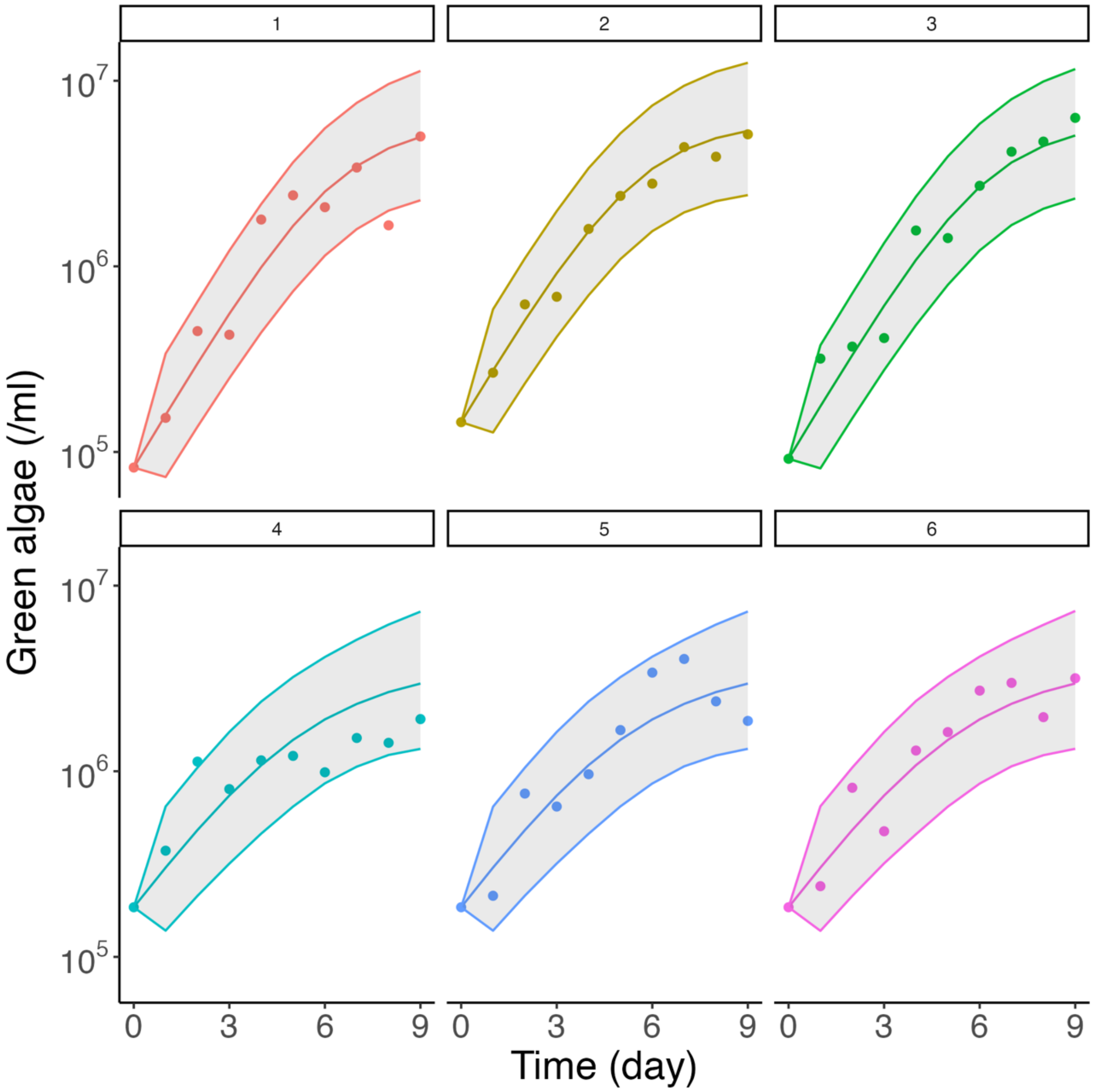


**Figure S4: Population dynamics of *Chlamydomonas sphaeroides* in mono-culture.**
The population dynamics of *Chlamydomonas sphaeroides* with (the top three panels) and without (the bottom three panels) salinity stress, with three replicates in each condition. The dots represent the mean densities of the green algae from three measurements, the shaded areas show the 95% CI, and the middle-colored lines indicate the dynamics with the median values from the posterior distribution.

### S1.3: Estimation of the rotifer parameters

Next, we estimated the attack rate, conversion efficiency, and mortality rates of rotifers in the absence or presence of salinity stress assuming that the conversion efficiency is not affected by salt. Since the mortality rate is expected to be independent of the prey species, we employed the following two-step approach. First, we fitted our model to the rotifer-*Chlorella* co-culture experiments to estimate the mortality rates with and without salinity stress. Then, we used the posterior distributions of the mortality rates with and without salinity stress as the *prior distributions* to fit the rotifer-*Chlamydomonas* co-culture experiments. The rationale is that the attack rate on *Chlorella vulgaris* would not change under salinity stress because *Chlorella* did not form clumps (Fig. 2). Thus, the salinity impact on the mortality rate would be more clearly inferred from the co-culture with *Chlorella vulgaris* than with *Chlamydomonas sphaeroides*.

Because we were not sure of the scales of the attack rate on *Chlorella vulgaris* and the mortality rate under salinity stress, we transformed them into log scales and set the prior distributions as following normal distributions:

$$\log m_j \sim N\,(\log 0.1\,, 1^2),$$
$$\log a_{11} = \log a_{12} \sim N\,(\log 1\,, 0.5^2).$$

Because the previous study indicates that the mortality rate of the rotifers without salinity stress is 0.055/day (Becks et al. 2010), we set the prior distribution of $m_j$ so that its mean is 0.16 and 95% CI lies between 0.014 and 0.71. The prior distribution of $a_{11} = a_{12}$ ($\equiv a_1$) was set so that its mean is 1.13 and 95% CI lies between 0.27 and 2.66 ml / rotifers / day.

The prior distribution of the conversion efficiency was inferred from the result of Becks et al. (2010). They showed the conversion efficiency from *Chlamydomonas reinharditii* to the rotifer was 170 rotifers / $10^6$ algal cells = $1.7 \times 10^3$ rotifers / $10^7$ algal cells. Because the cell volume of *Chlamydomonas reinharditii* is ca. $2^3$~$2.5^3$ times larger than *Chlorella vulgaris* and we scaled the density of *Chlorella vulgaris* by dividing by $10^8$, we used the following prior distribution:

$$c_1 \sim N(1.5, 0.2^2).$$

We ran 50,000 MCMC iterations for each estimation with 5,000 warm-ups and four chains. The posterior distributions of the parameter values and the fitting results are shown in Figs. S5-S7. We found that salinity stress indeed increased the rotifer mortality rate (i.e., the direct effect: Fig. S7).

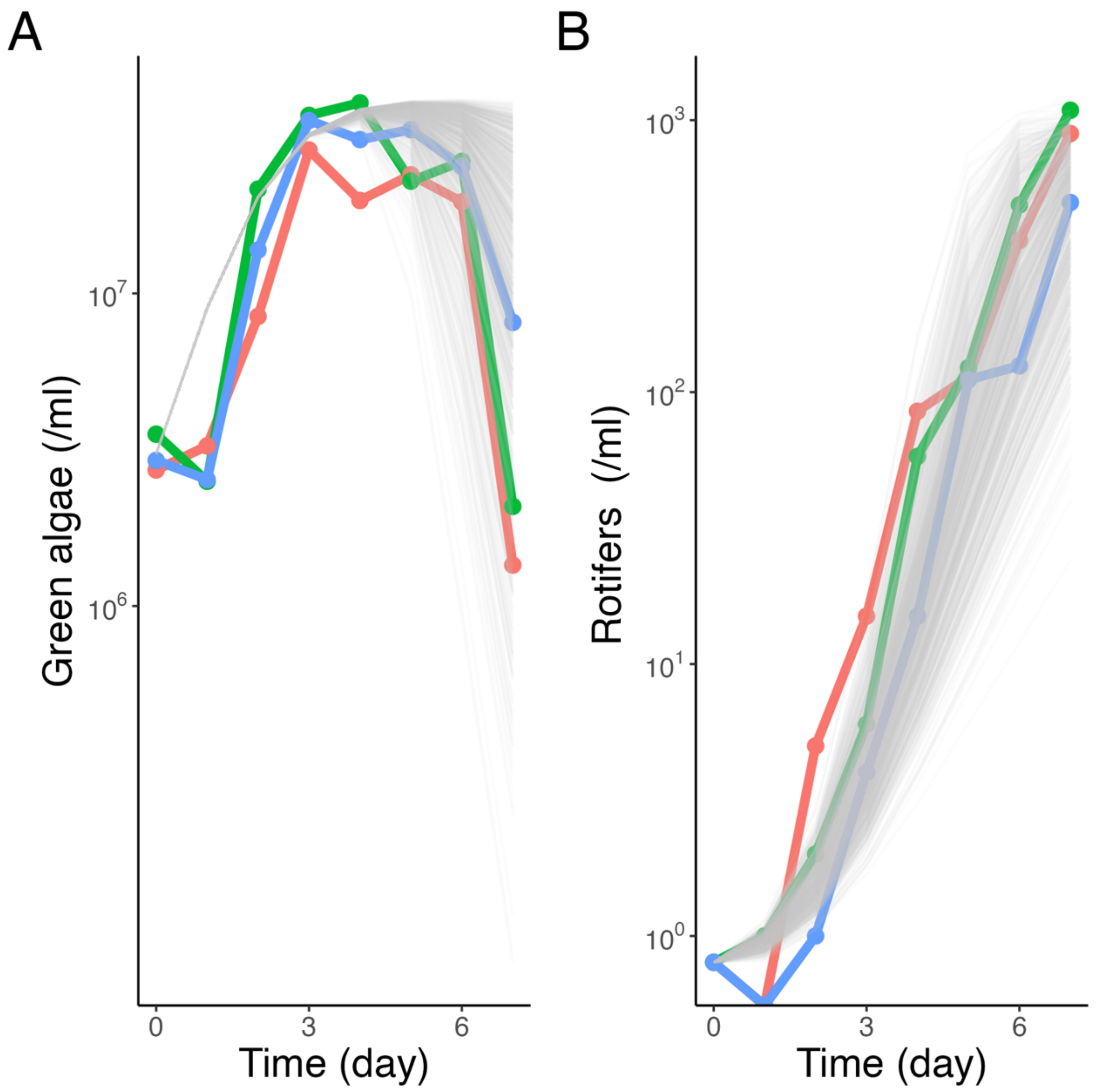


**Figure S5: Population dynamics of *Chlorella vulgaris* and rotifers without salinity stress.** Fitting to the empirical dynamics of the green algae (A) and the rotifers (B). Each color corresponds to a different replicate. The gray lines represent 1,000 simulations with sampling parameter values from the posterior distributions (Fig. S7).

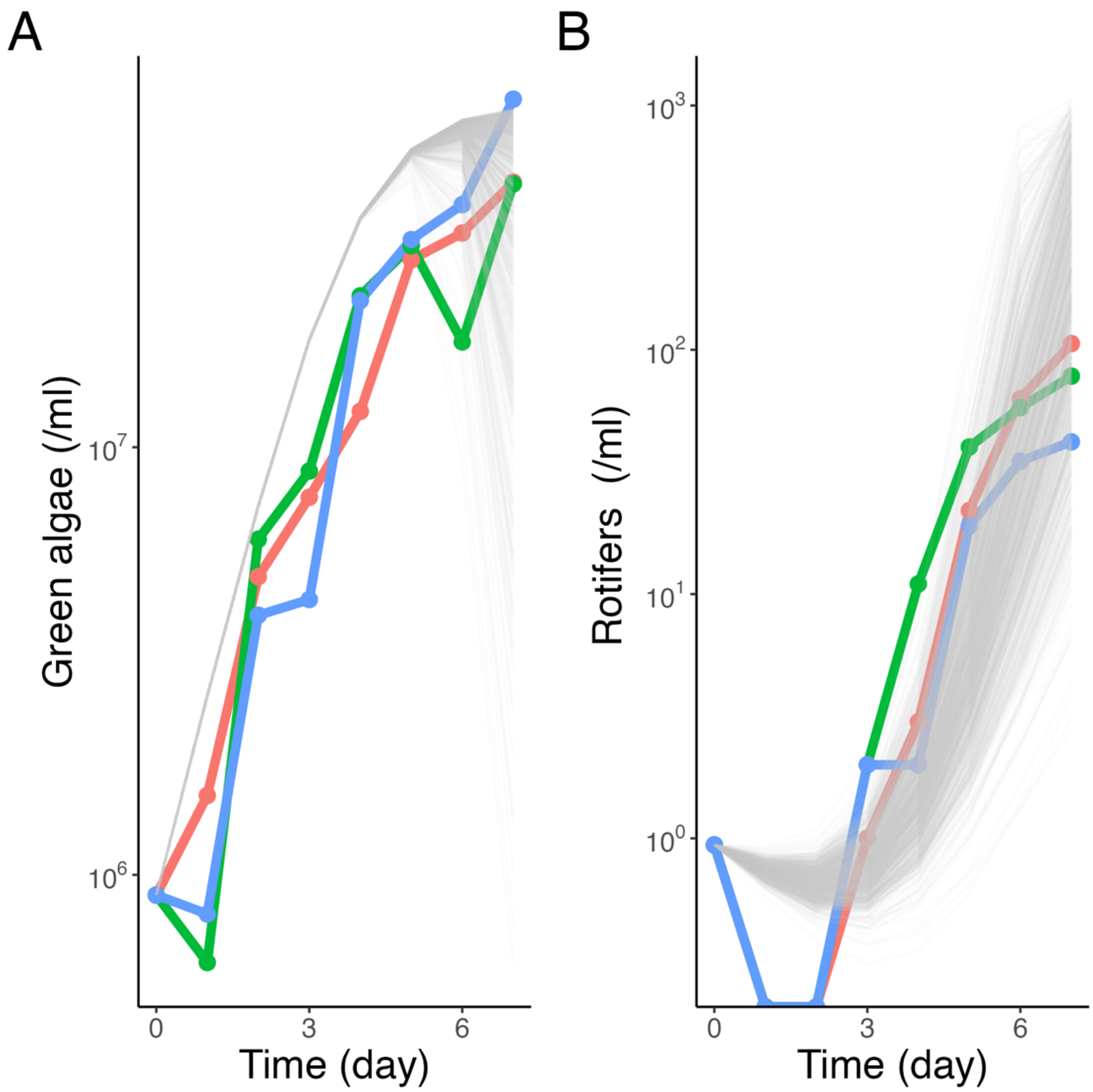


**Figure S6: Population dynamics of *Chlorella vulgaris* and rotifers with salinity stress.** Fitting to the empirical dynamics of the green algae (A) and the rotifers (B). Each color corresponds to a different replicate. The gray lines represent 1,000 simulations with sampling parameter values from the posterior distributions (Fig. S7).

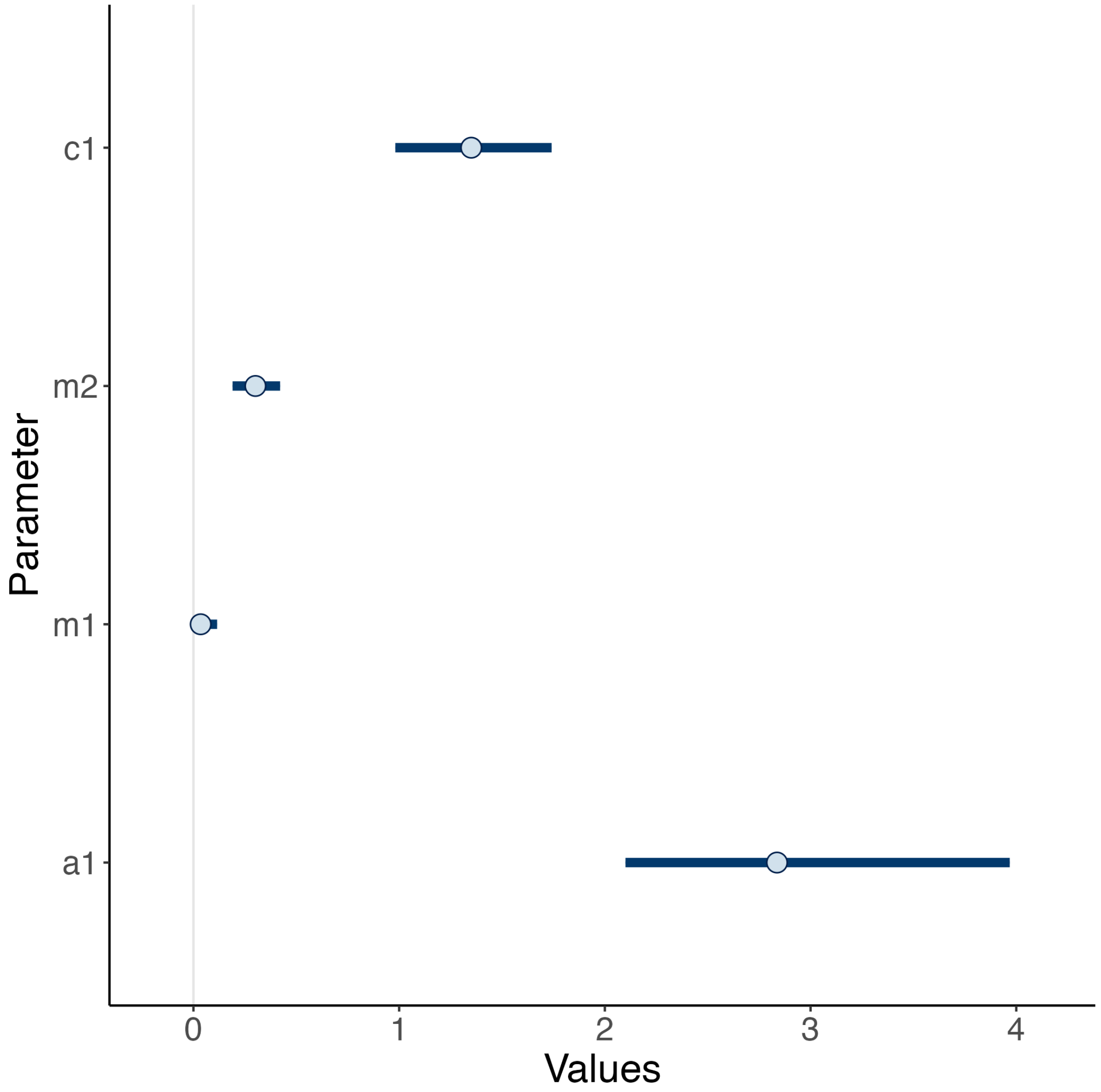


**Figure S7: The posterior distributions of the parameters after fitting to the co-culture data of *Chlorella vulgaris* and rotifers.**

The posterior distributions of the attack rate on *Chlorella vulgaris* ($a_1 = a_{11} = a_{12}$), the mortality rates of the rotifer in the absence ($m_1$) or presence ($m_2$) of salinity stress, and the conversion efficiency ($c_1$) are shown. The dots indicate the median values, while the horizontal lines represent the 95% CIs.

Finally, we fitted our model to the data of rotifer-*Chlamydomonas* co-culture experiments. We set the prior distributions as follows:

$$a_{2j} \sim N\,(0.6, 0.2^2),$$
$$c_2 \sim N\,(1.7, 0.2^2),$$
$$\log m_j \sim N\left(\log \mu_j, \log s_j^2\right),$$

where $\log\mu_j$ and $\log s_j^2$ are the mean and standard deviation of the posterior probability distribution of $\log m_j$ after fitting to the co-culture experiment data of the rotifers with *Chlorella vulgaris* (Fig. S7). The mean of the prior distributions of the attack rates was assumed to be smaller than that on *Chlorella vulgaris*, and we determined the mean value for MCMC to successfully converge. The prior distribution of the conversion efficiency was inferred from the previous study (Becks et al. 2010) by assuming that the conversion efficiency of *Chlamydomonas sphaeroides* is similar to that of *Chlamydomonas reinharditii*. The prior distributions of the mortality rates were generated from their posterior distribution after fitting to the rotifer-*Chlorella* co-culture experiment data.

We ran 50,000 MCMC iterations for each estimation with 5,000 warm-ups and four chains. The posterior distributions of the parameter values and the fitting results are shown in Figs. S8-S10. The posterior distributions show that the attack and mortality rates changed due to salinity stress (see also Fig. 4): we found that salinity stress indeed increased the rotifer mortality rate (i.e., the direct effect) and it also decreased the attack rate probably due to algal clumping (i.e., the trait-mediated indirect effect: Fig. S10).

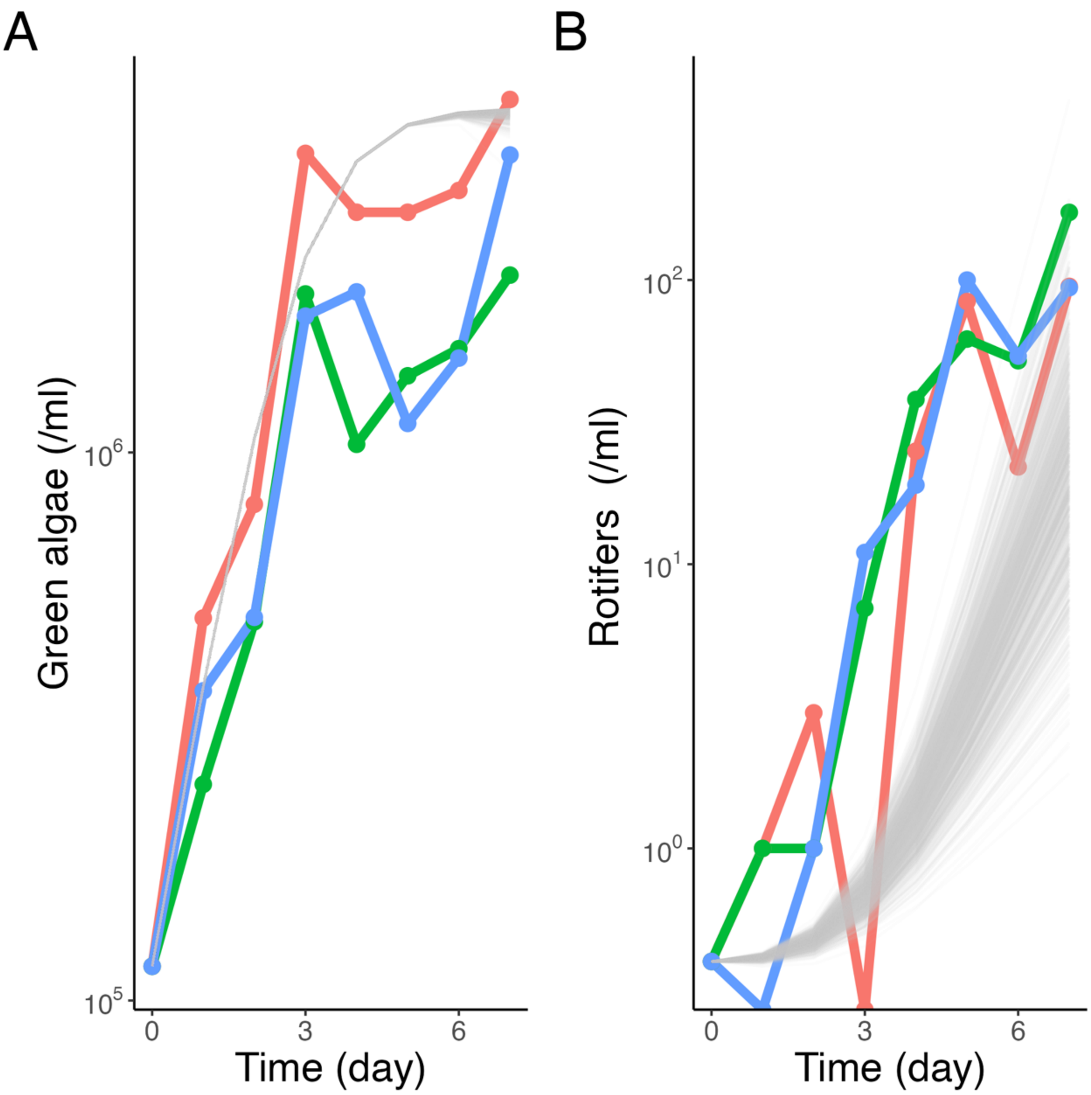


**Figure S8: Population dynamics of *Chlamydomonas sphaeroides* and rotifers without salinity stress.**

Fitting to the empirical dynamics of the green algae (A) and the rotifers (B). Each color corresponds to a different replicate. The gray lines represent 1,000 simulations with sampling parameter values from the posterior distributions (Fig. S10).

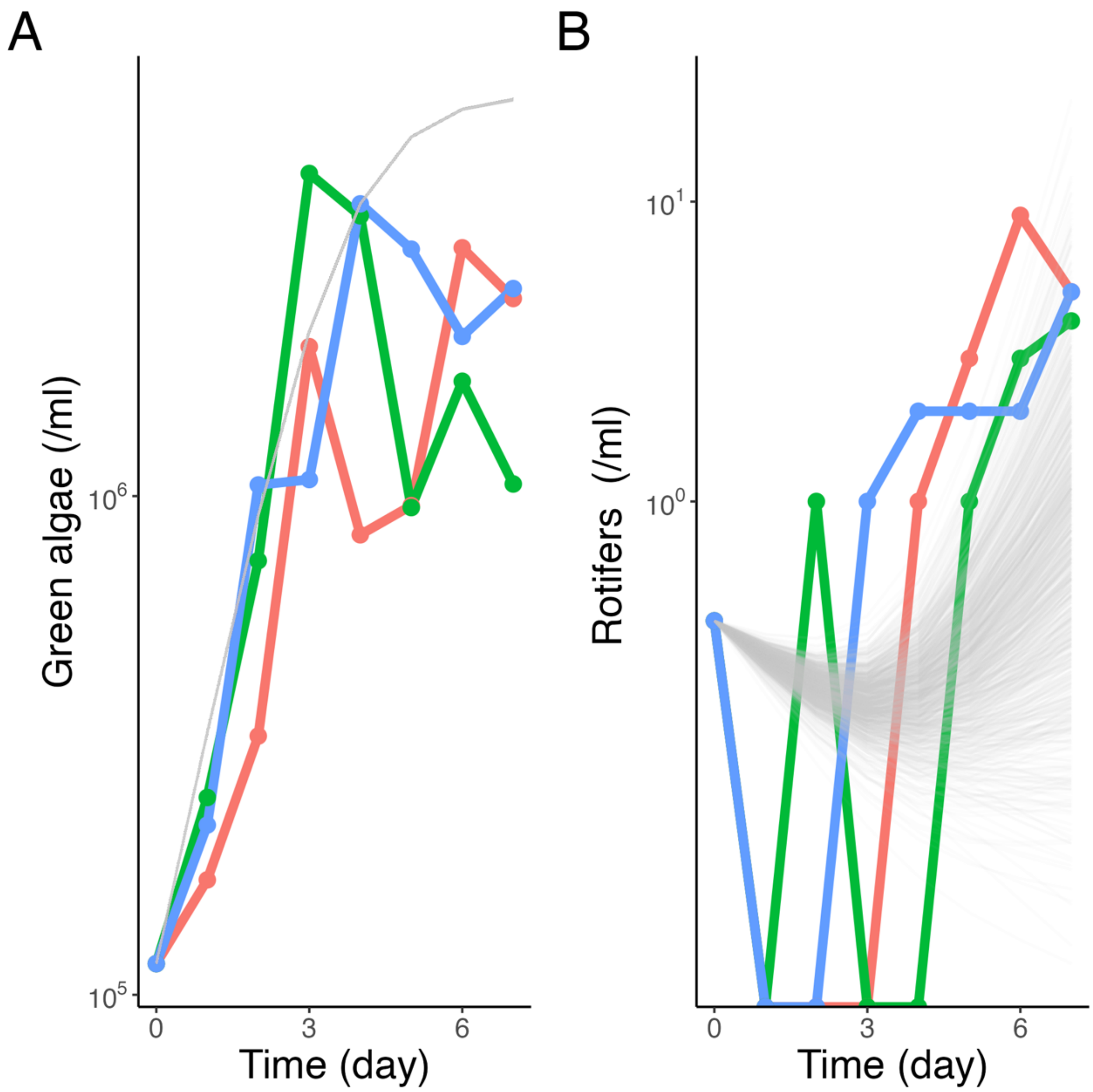


**Figure S9: Population dynamics of *Chlamydomonas sphaeroides* and rotifers with salinity stress.**

Fitting to the empirical dynamics of the green algae (A) and the rotifers (B). Each color corresponds to a different replicate. The gray lines represent 1,000 simulations with sampling parameter values from the posterior distributions (Fig. S10).

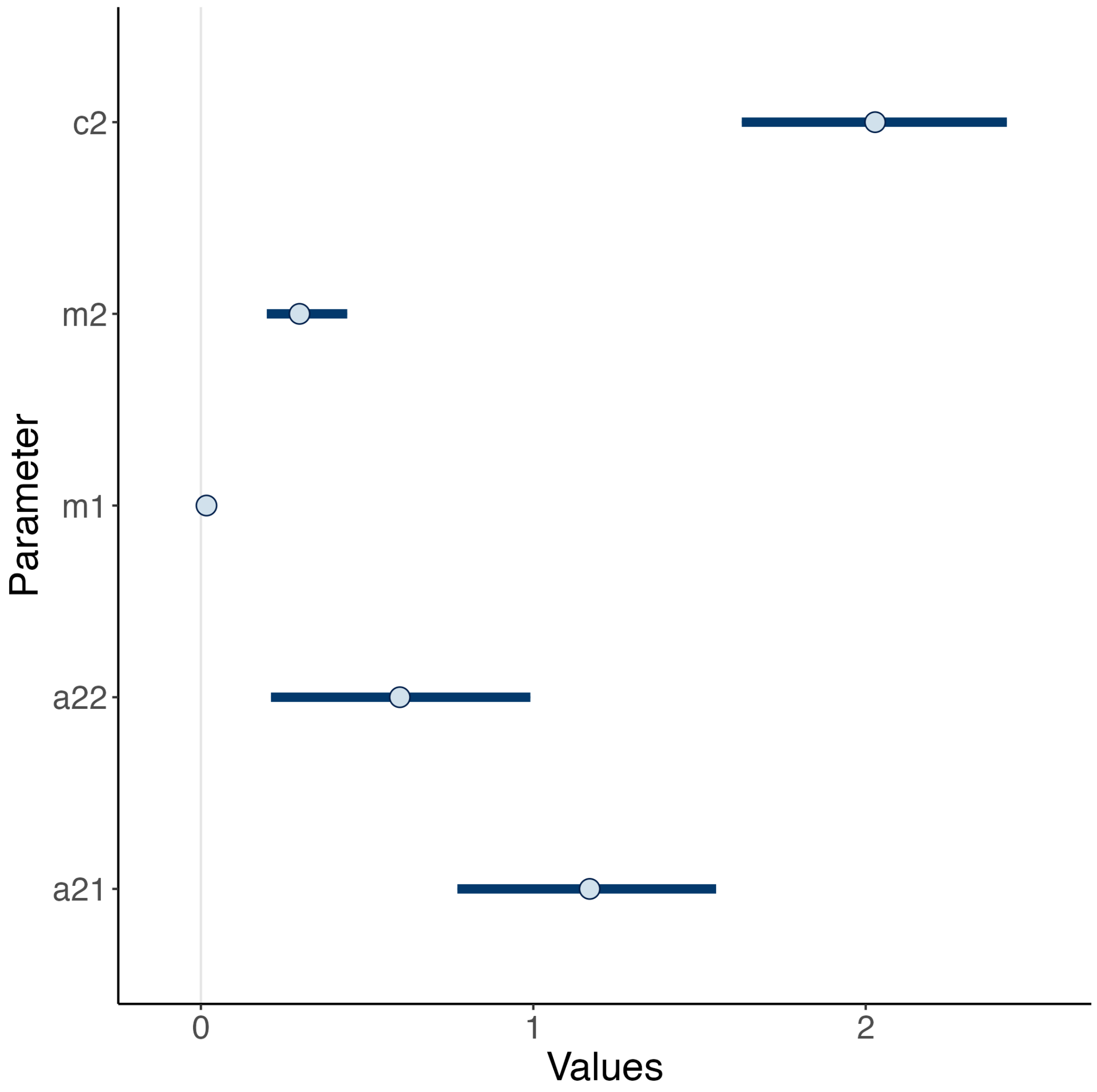


**Figure S10: The posterior distributions of the parameters after fitting to the co-culture data of *Chlamydomonas sphaeroides* and rotifers.**

The posterior distributions of the attack rate on *Chlamydomonas sphaeroide*s in the absence ($a_{21}$) or presence ($a_{22}$) of salinity stress, the mortality rates of the rotifer in the absence ($m_1$) or presence ($m_2$) of salinity stress, and the conversion efficiency ($c_2$) are shown. The dots indicate the median values, while the horizontal lines represent the 95% CIs.

### S2: Results of the experimental evolution of *Chlamydomonas sphaeroides*

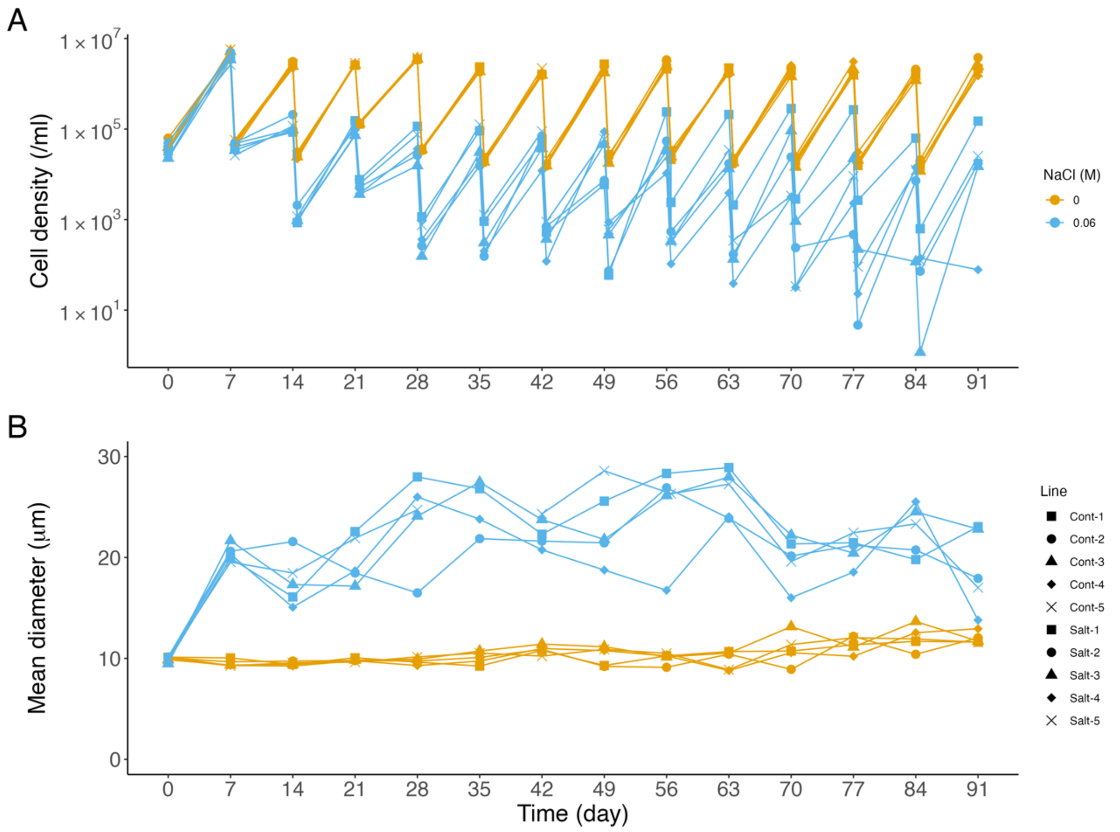


**Figure S11: Population and mean diameter dynamics of *Chlamydomonas sphaeroides*.** The 91-day experimental results of population dynamics (A) and mean diameter (B) are shown. The blue lines represent the dynamics under 0.06M NaCl, while the orange lines show the control ones (i.e., 0M NaCl). The shape of each marker indicates each replicate.

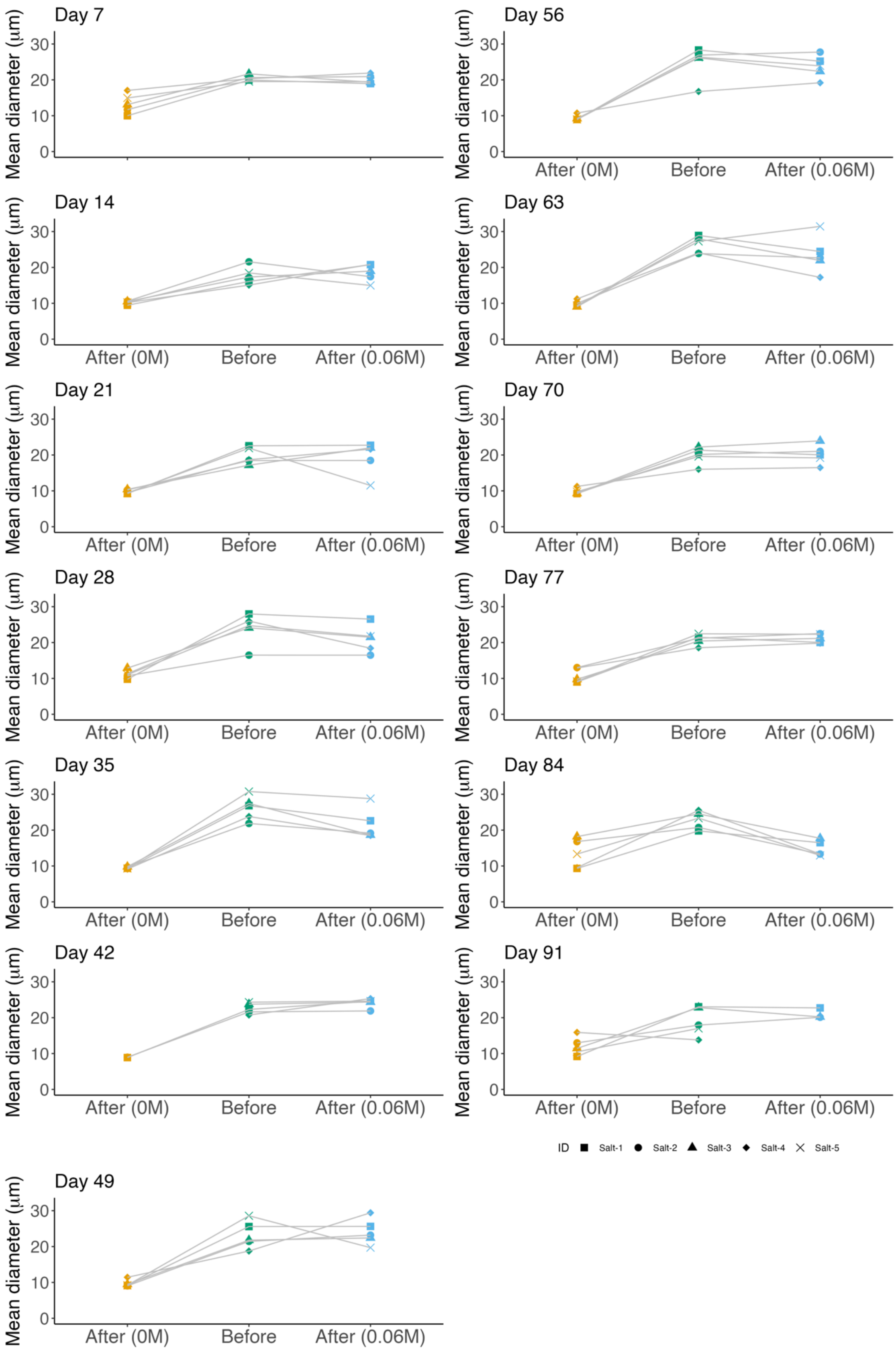
Day 7
Day 14
Day 21
Day 28
Day 35
Day 42
Day 49
Day 56
Day 63
Day 70
Day 77
Day 84
Day 91
Mean diameter (µm)
0
10
20
30
After (0M)
Before
After (0.06M)
ID
Salt-1
Salt-2
Salt-3
Salt-4
Salt-5

**Figure S12: Mean diameter of particles during the 91-day experiment.**
We measured the mean diameter of particles under salinity stress every week during the 91-day experiment (green dots). A part of each culture was then transferred to a fresh C medium with either 0M (orange) or 0.06M (blue) NaCl, and cultivated for six hours. If the mean diameter of particles in 0M NaCl was as high as before transfer to the fresh media, the green algae lose phenotypic plasticity, and multicellular clumps remain without salinity stress.
